# Quantification of Spin Fluctuations during Stem Cell Differentiation Using Diamond Nanopillars

**DOI:** 10.64898/2026.11.16.767033

**Authors:** Abdul Nasir Kuzhiyan Thadathil, Yoko Shigeno, Hitoshi Niwa, Hitoshi Ishiwata

## Abstract

Quantitative detection of radical-associated magnetic activity in living cells remains challenging because variations between quantum sensors complicate direct comparisons of intracellular environments. Here, we demonstrate a label-free quantum sensing platform based on substrate-fixed diamond nanopillars containing individually addressable nitrogen-vacancy (NV) centers. The fixed architecture enables one-to-one measurements using identical NV sensors in the presence and absence of cells, minimizing sensor-to-sensor variability and allowing reliable assessment of cellular contributions to spin decoherence. Cellular compatibility assays supported preserved metabolic activity, DNA synthesis, and membrane integrity without detectable apoptotic activation. Intracellular NV centers retained approximately 10% optically detected magnetic resonance contrast, 20% Rabi contrast, and a Hahn-echo coherence time (T₂) of approximately 3 µs. Carr–Purcell–Meiboom–Gill dynamical decoupling extended intracellular T₂ to 25.8 ± 4.90 µs. We applied this platform to embryonic stem (ES) cells undergoing doxycycline-induced differentiation toward the trophectoderm lineage. Identical NV sensors exhibited consistent coherence recovery following removal of differentiation-induced cells, whereas undifferentiated controls showed no systematic recovery. Calibration using the stable radical TEMPOL, supported by magnetic-noise modeling and kinetic Monte Carlo simulations, enabled numerical evaluation of local magnetic responses as model-dependent TEMPOL-equivalent spin densities of 0.084–0.567 spins/nm³ across different sensing sites. These findings establish a quantitative approach to investigating intracellular physicochemical changes during stem-cell differentiation and demonstrate the potential of quantum sensing to study biological processes relevant to tissue development and regenerative medicine.

## Introduction

Embryonic stem (ES) cells can differentiate into specialized cell types and provide an important model for understanding how cellular changes contribute to tissue development and biological function. The ability to monitor stem-cell differentiation is also essential for advancing regenerative medicine, where quantitative evaluation of cellular processes is needed to understand and control the acquisition of specialized functions. Reactive oxygen species (ROS) and free radicals participate in stem-cell proliferation, differentiation, and cellular signaling [1–6]. Although excessive ROS can cause oxidative damage, controlled radical production contributes to normal cellular functions. Conventional fluorescence-based ROS probes enable visualization of intracellular oxidative activity but are affected by variations in dye uptake, photobleaching, and local chemical conditions [7–10]. Quantitative, label-free measurements of radical-associated activity could therefore provide new insights into the physicochemical changes accompanying stem-cell differentiation.

Nitrogen-vacancy (NV) centers in diamond offer a promising approach for detecting local spin fluctuations associated with nearby paramagnetic species through spin relaxation and coherence measurements. Previous intracellular NV studies have investigated free-radical-associated responses in various biological systems [11–18], including functional changes in immune cells and cardiac fibroblasts [19,20]. These studies demonstrate the potential of quantum sensing for investigating cellular functions. However, extending this approach to the differentiation of pluripotent stem cells would enable quantum measurements of biological processes associated with the formation of specialized cell types and, ultimately, tissue-level functions. A major challenge is achieving reliable quantitative measurements in living cells. Freely dispersed nanodiamonds vary in position, orientation, and intrinsic spin properties, making it difficult to distinguish genuine changes in the cellular magnetic environment from sensor-to-sensor variability [21–23]. Previous studies have demonstrated the difficulty of maintaining long spin coherence times inside living cells, even with nanodiamonds exhibiting relatively long coherence times outside cells [22].

Reliable quantitative intracellular quantum sensing therefore benefits from one-to-one comparisons using the same quantum sensor in the presence and absence of cells. Substrate-fixed diamond nanopillar arrays provide individually addressable NV centers at precisely defined positions [24,25]. Their fixed geometry allows the same NV sensor to be repeatedly identified and measured with cells present and again following cell removal, minimizing uncertainties arising from differences between individual sensors. Furthermore, the nanopillar architecture supports coherent quantum measurements inside living cells while maintaining the cellular functions assessed experimentally. Combined with calibration using known paramagnetic radicals, this approach provides a quantitative framework for converting changes in NV spin coherence into numerical estimates of local magnetic activity, rather than relying solely on qualitative comparisons.

Here, we demonstrate a substrate-fixed diamond nanopillar quantum sensing platform for investigating embryonic stem-cell differentiation induced by doxycycline-mediated Oct3/4 repression toward the trophectoderm lineage [26,27]. The nanopillars maintain coherent NV spin manipulation inside living cells, including a Hahn-echo T₂ of approximately 3 µs and a CPMG-enhanced coherence time of 25.8 ± 4.90 µs. Using individually tracked sensors, we performed direct one-to-one comparisons before and after cell removal and detected spin fluctuations associated with differentiation. Reference measurements using the stable radical TEMPOL, together with Gaussian magnetic-noise modeling and complementary kinetic Monte Carlo simulations, enabled quantitative comparison of local magnetic responses as model-dependent TEMPOL-equivalent spin densities of 0.084–0.567 spins/nm³ across different locations. By extending intracellular quantum sensing from functional changes in differentiated cells to the differentiation of pluripotent stem cells, this work establishes a strategy for quantitatively investigating cellular processes relevant to tissue formation and regenerative medicine.

## Results and discussions

### Diamond Nanopillar-Based Quantum Sensing Platform for Intracellular Measurements

We developed a substrate-fixed diamond nanopillar platform containing nitrogen-vacancy (NV) centers for one-to-one measurements of local spin fluctuations in cellular environments. Figure 1(A) illustrates cells cultured on diamond nanopillar arrays with near-surface NV centers. Individual pillars are addressed by optical excitation, fluorescence readout, and coherent microwave manipulation. The fixed grid permits each NV sensor to be identified and measured under cell-present conditions and remeasured after removal of the same cellular environment. In contrast to comparisons between freely dispersed nanodiamonds, this paired design minimizes differences arising from NV depth, orientation, surface properties, and initial coherence among sensors. Nanopillars not covered by cells also permit contemporaneous extracellular measurements, although the principal one-to-one comparison uses the same site before and after cell removal. ROS-reactive fluorescent probes are not required for the NV magnetic measurement. Scanning electron microscopy (SEM) images of a representative nanopillar and the array are shown in Figures 1(B) and 1(C), respectively. The nanopillars are 300 nm in diameter and 3.2 µm in height, with a center-to-center spacing of 15 µm and near-surface NV centers approximately 10 nm below the diamond surface. This approximate depth, used in the subsequent magnetic-noise calculations, was based on experimental NV-depth measurements reported for comparable 3 keV nitrogen implantation conditions [28].

**Figure 1.**
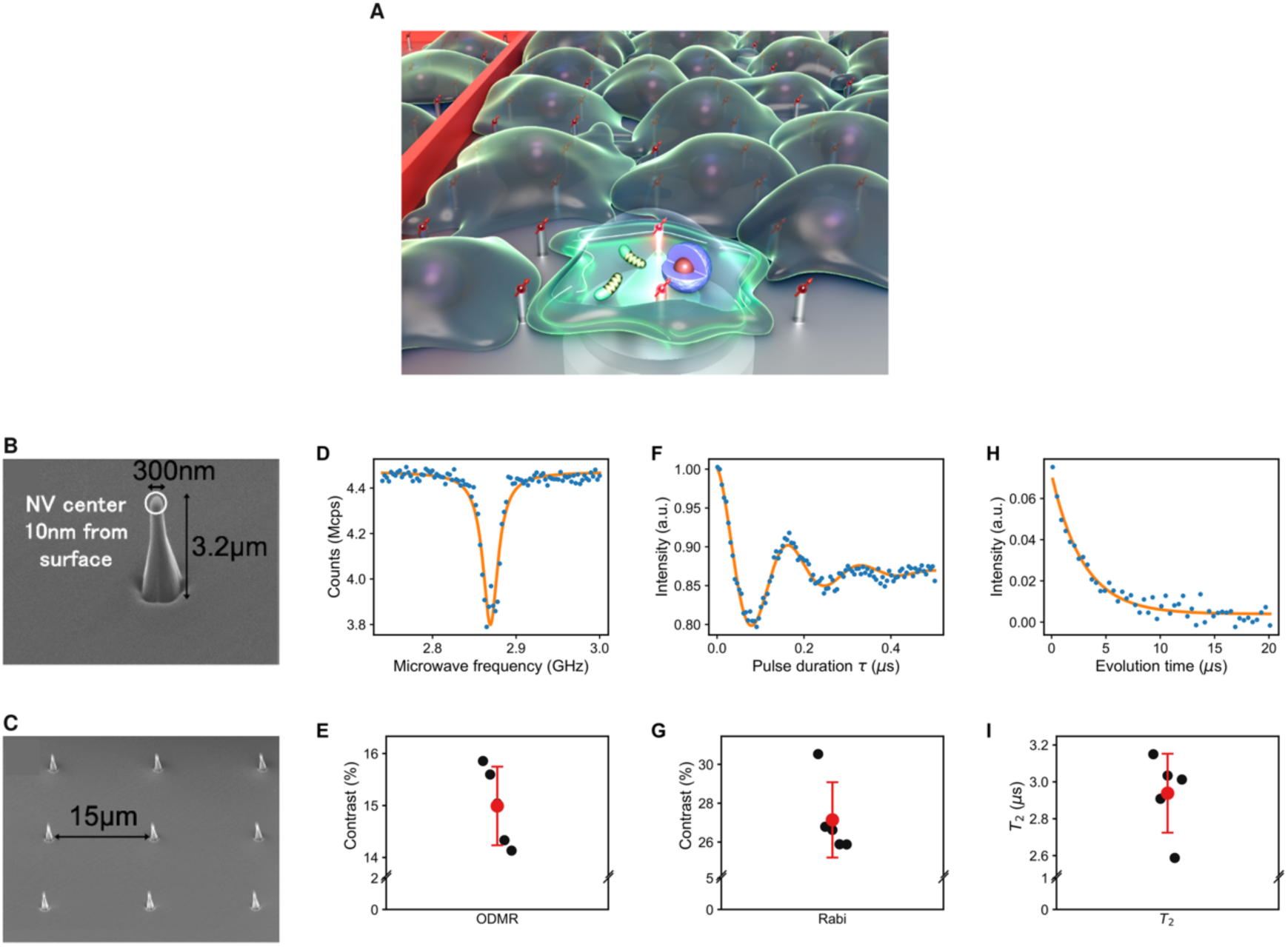
Characterization of substrate-fixed diamond nanopillar NV quantum sensors. (A) Schematic illustration of intracellular quantum sensing using NV-containing diamond nanopillars. (B) Scanning electron microscopy (SEM) image of an individual diamond nanopillar. (C) SEM image of a diamond nanopillar array. (D, E) Representative optically detected magnetic resonance (ODMR) spectrum and ODMR contrast measured across five nanopillars. (F, G) Representative Rabi oscillations and corresponding Rabi contrast measurements. (H, I) Representative Hahn spin-echo coherence decay and coherence times (T₂) measured across five nanopillars.

The quantum sensing performance of the diamond nanopillars was characterized using optically detected magnetic resonance (ODMR) spectroscopy, Rabi oscillations, and Hahn spin-echo measurements. Representative results are shown in Figures 1(D), 1(F), and 1(H), and the measurements across five nanopillars are summarized in Figures 1(E), 1(G), and 1(I). The mean ODMR and Rabi contrasts were 14.9 ± 0.76% and 27.1 ± 1.94%, respectively. Single-exponential fits to the Hahn-echo decays yielded a mean coherence time (T₂) of 2.94 ± 0.21 µs (Figure 1(I)). Beyond providing optical contrast and coherent microwave control, the fixed array design enables the baseline response of each individual sensor to be measured and distinguished from its subsequent response to the cellular environment. This reproducible sensor identity is central to the quantitative comparisons described below.

### Cellular Compatibility and Intracellular Quantum Coherence of Diamond Nanopillar Sensors

To evaluate the suitability of diamond nanopillars for intracellular quantum sensing, we first assessed their cellular compatibility using Hepa1-6 cells through fluorescence-based measurements of metabolic activity, apoptosis, DNA synthesis, and plasma membrane integrity (Figures 2(A–H)). Cellular metabolic activity was evaluated using the alamarBlue Cell Viability Reagent (Invitrogen, DAL1100), which detects the reduction of resazurin to fluorescent resorufin by metabolically active cells. The assay was performed using the manufacturer-recommended working conditions, with resorufin fluorescence monitored at approximately 560 nm excitation and 590 nm emission. Cells cultured on diamond nanopillar arrays exhibited clear fluorescence, indicating preserved metabolic activity (Figure 2(A)), whereas dead-cell controls showed no detectable signal (Figure 2(B)). Apoptotic activity was evaluated using the CellEvent Caspase-3/7 Green Detection Reagent (Thermo Fisher Scientific, C10723), a fluorogenic substrate that produces green fluorescence upon cleavage by activated caspase-3/7, with representative excitation and emission wavelengths of approximately 502 and 530 nm, respectively. Cells cultured on nanopillars exhibited no detectable caspase-3/7 fluorescence (Figure 2(C)), in contrast to the strong fluorescence observed in apoptosis-positive controls (Figure 2(D)). These results indicate that cell–nanopillar interactions did not produce detectable impairment of metabolic activity or apoptotic activation under the experimental conditions.

**Figure 2.**
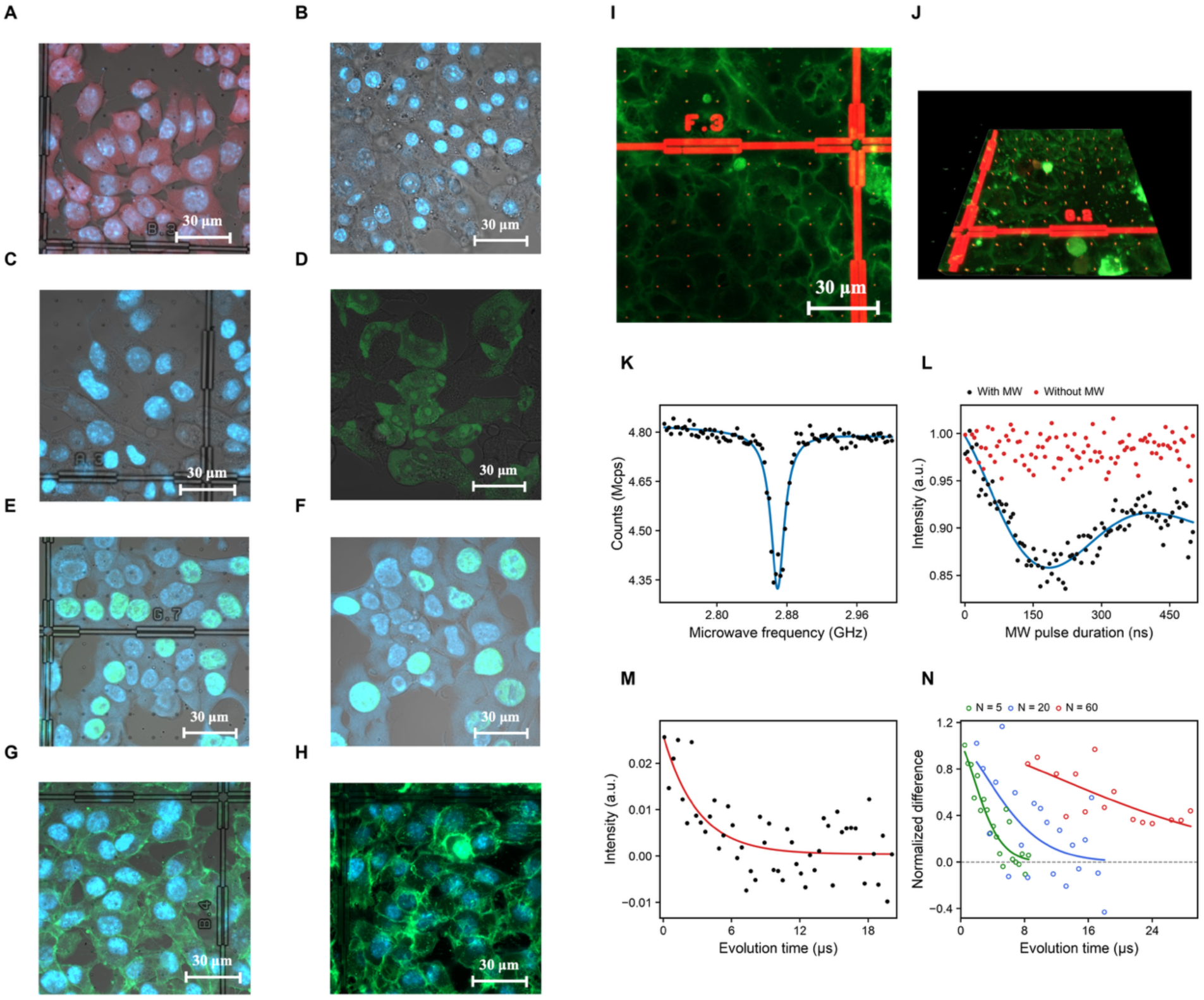
Cellular compatibility and intracellular quantum coherence of diamond nanopillar NV sensors. (A, B) Metabolic activity of Hepa1-6 cells assessed using the alamarBlue assay under living-cell (A) and dead-cell control (B) conditions. (C, D) Caspase-3/7 fluorescence assays showing nanopillar-associated cells (C) and apoptosis-positive controls (D). (E, F) EdU incorporation in cells cultured on diamond nanopillar arrays (E) and outside the nanopillar regions (F). (G, H) Fluorescence images of plasma membrane integrity and nuclear morphology in cells cultured on nanopillars (G) and dead-cell controls (H). (I, J) Confocal fluorescence images showing diamond nanopillars associated with living cells. (K) Representative intracellular optically detected magnetic resonance (ODMR) spectrum. (L) Intracellular Rabi oscillations demonstrating coherent NV spin manipulation. (M) Intracellular Hahn spin-echo coherence decay. (N) Carr–Purcell–Meiboom–Gill (CPMG) coherence decay curves for different refocusing pulse numbers (N = 5, 20, and 60), demonstrating extended spin coherence inside living cells.

DNA synthesis was further examined using the Click-iT EdU Cell Proliferation Kit (Thermo Fisher Scientific, C10337). Following EdU incorporation into newly synthesized DNA during S phase, cells were fixed, permeabilized, and fluorescently labeled using the click-chemistry reaction according to the manufacturer’s protocol. Cell nuclei were counterstained with Hoechst dye, which exhibits characteristic blue fluorescence (excitation approximately 350 nm; emission approximately 461 nm). EdU-positive cells were observed both on the nanopillar arrays (Figure 2(E)) and outside the nanopillar regions (Figure 2(F)), demonstrating that DNA synthesis was maintained in cells interacting with the nanopillars. Plasma membrane morphology and integrity were evaluated using CellMask Plasma Membrane Stain (Invitrogen, C37608), NucBlue Live, and NucGreen Dead (Invitrogen, R37609). CellMask delineated the plasma membrane, NucBlue labeled nuclei, and NucGreen Dead identified cells with compromised plasma membrane integrity through nuclear fluorescence. Cells cultured on nanopillars exhibited well-defined plasma membrane and nuclear morphology without detectable NucGreen nuclear fluorescence (Figure 2(G)), whereas dead-cell controls showed NucGreen-positive nuclei (Figure 2(H)). Collectively, these complementary assays indicate that the diamond nanopillar platform preserves detectable metabolic activity, DNA synthesis, and plasma membrane integrity without evidence of apoptotic activation, supporting its cellular compatibility under the investigated conditions.

We next investigated whether NV centers within the diamond nanopillars retained their optical spin readout and coherent quantum control capabilities in living cells. Confocal laser-scanning microscopy identified nanopillars located within the cellular environment (Figures 2(I) and 2(J)), enabling site-specific optical interrogation of individual NV-containing nanopillars. The NV centers were optically excited using a 532 nm laser, and their spin-dependent fluorescence was collected in the red emission region using an appropriate optical detection pathway. Microwave excitation was applied near the NV electron-spin resonance frequencies to perform optically detected magnetic resonance (ODMR) spectroscopy and pulsed spin measurements. Intracellular ODMR measurements yielded a contrast of approximately 10% (Figure 2(K)), while coherent microwave manipulation produced Rabi oscillations with a contrast of approximately 20% (Figure 2(L)). These results demonstrate efficient spin-dependent optical readout and coherent spin manipulation in the cellular environment.

Spin coherence was further characterized by Hahn spin-echo measurements, yielding T₂ of approximately 3 µs inside living cells (Figure 2(M)). We then applied Carr–Purcell–Meiboom–Gill (CPMG) dynamical decoupling to suppress dephasing from environmental spin fluctuations over the frequencies addressed by the pulse sequence. The coherence time increased with the number of refocusing pulses, reaching T₂ = 3.48 ± 0.68 µs (N = 5), 7.04 ± 3.75 µs (N = 20), and 25.8 ± 4.90 µs (N = 60) (Figure 2(N)). The N = 60 value corresponds to approximately 8.6-fold extension relative to the intracellular Hahn-echo T₂ of approximately 3 µs. For context, Oshimi et al. reported a Hahn-echo T₂ of 3.2 ± 1.2 µs for free nanodiamonds outside cells but an apparent intracellular spin-echo decay time of approximately 0.135 µs, which was affected by particle motion and orientation [22]. Although the device configurations and measurement conditions differ, our results show that substrate-fixed nanopillars maintain coherent control inside living cells and permit effective pulsed CPMG decoupling with exceptionally long intracellular coherence.

Together, the cellular compatibility assays and intracellular spin measurements support the use of substrate-fixed nanopillars for label-free quantum biosensing under the conditions examined. The alamarBlue, caspase-3/7, EdU, and membrane-integrity assays provide complementary evidence that metabolic activity, DNA synthesis, and membrane integrity are maintained without detectable apoptotic activation. Fluorescent reagents were used for these independent biological assessments, whereas NV-based readout of spin fluctuations does not require ROS-reactive probes. Importantly, the same fixed sensing locations can be revisited for paired measurements with cells present and after cell removal. The combination of cellular compatibility, coherent spin control, and repeatable sensor identity provides a basis for investigating local paramagnetic activity during stem-cell differentiation.

### Label-Free Detection of Differentiation-Associated Spin fluctuations in Embryonic Stem Cells

Having established cellular compatibility and coherent intracellular operation, we next investigated magnetic changes associated with the differentiation of embryonic stem (ES) cells. Previous NV-center studies have primarily examined free-radical-associated responses of mature cell types, including macrophages and cardiac fibroblasts [19,20]. Here, we extend quantum sensing to pluripotent ES-cell differentiation, an experimentally accessible model for understanding how local cellular changes accompany the development of specialized functions. Differentiation toward the trophectoderm lineage was induced using a Tet-Off system in which doxycycline (DOX) suppresses Oct3/4, a transcription factor required to maintain pluripotency [26,27]. Representative images of ES cells under untreated (DOX−) and differentiation-inducing (DOX+) conditions are shown in Figures 3(A) and 3(B), respectively. This cellular model provides a first step toward applying quantitative quantum readouts to developmental processes of potential relevance to tissue function and regenerative medicine, without implying that tissue-level function was directly measured here.

**Figure 3.**
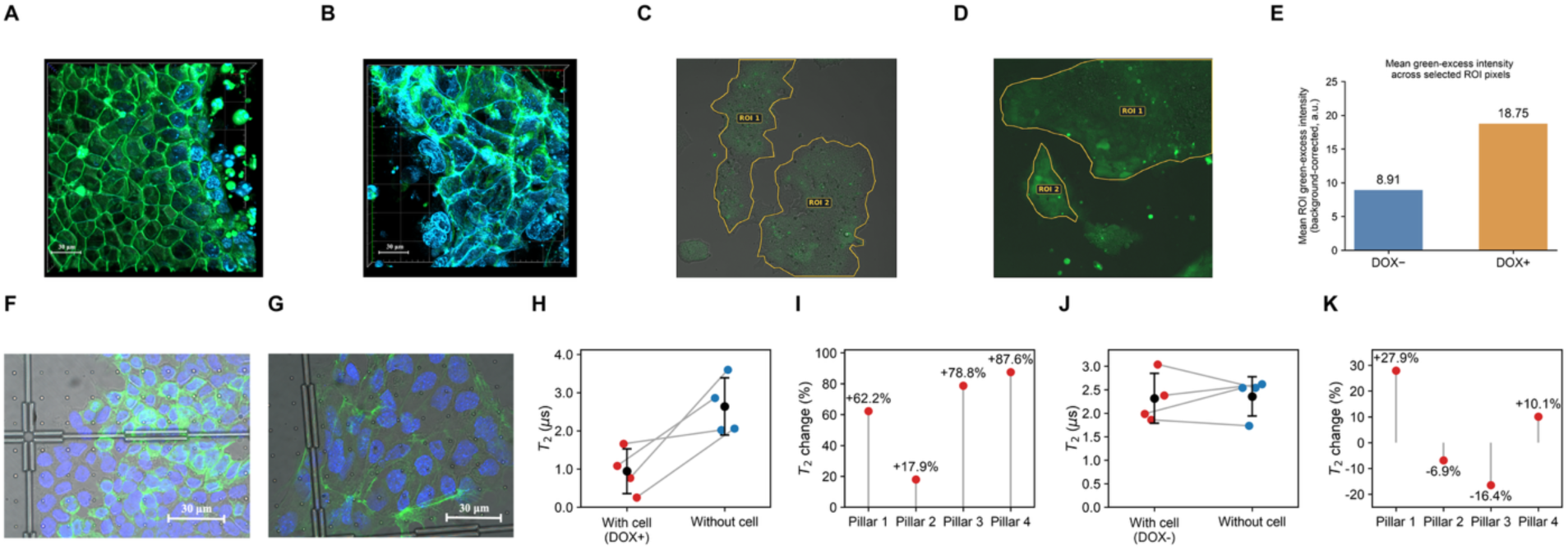
Label-free detection of differentiation-associated spin fluctuations in embryonic stem cells using diamond nanopillar NV sensors. (A, B) Confocal fluorescence images of embryonic stem (ES) cells under undifferentiated (DOX−, A) and differentiation-inducing (DOX+, B) conditions. (C, D) Oxidation-sensitive CM-H₂DCFDA fluorescence images of ES cells under DOX− (C) and DOX+ (D) conditions. (E) Comparison of background-corrected fluorescence intensities between DOX− and DOX+ cells. (F, G) Fluorescence images of GFP-labeled actin filaments and Hoechst-stained nuclei in DOX− (F) and DOX+ (G) cells, illustrating differentiation-associated morphological changes. (H) Paired Hahn spin-echo coherence times (T₂) measured using the same NV nanopillars in the presence of DOX-treated ES cells and after cell removal. (I) Corresponding percentage changes in T₂ for four individually identified nanopillars under DOX+ conditions. (J) Paired T₂ measurements for nanopillars associated with undifferentiated (DOX−) ES cells before and after cell removal. (K) Corresponding percentage changes in T₂ for four individually identified nanopillars under DOX− conditions. (L) Comparison of T₂ values measured at nanopillar locations classified as intracellular and extracellular under DOX+ conditions.

To evaluate changes in intracellular oxidative activity associated with differentiation induction, we performed fluorescence measurements using the oxidation-sensitive probe CM-H₂DCFDA. The fluorescence signal was analyzed within regions of interest (ROIs) selected to encompass cellular regions exhibiting green fluorescence (Figures 3(C) and 3(D)). DOX− ES cells exhibited relatively weak green fluorescence, whereas DOX+ cells showed substantially enhanced fluorescence. Quantitative analysis of the background-corrected mean green fluorescence intensity within the selected ROIs yielded values of 8.91 and 18.75 a.u. for DOX− and DOX+ cells, respectively (Figure 3(E)), corresponding to an approximately 2.1-fold increase under differentiation-inducing conditions. Although this comparison was based on representative images, the enhanced oxidation-sensitive fluorescence is consistent with increased intracellular oxidative activity following DOX treatment. We further examined cellular morphology and cytoskeletal organization by imaging GFP-labeled actin filaments and Hoechst-stained nuclei. Comparison of DOX− and DOX+ cells revealed differences in cellular morphology and actin organization (Figures 3(F) and 3(G)), providing complementary evidence of cellular responses to differentiation induction.

We then examined whether differentiation-associated cellular changes altered the local magnetic environment detected by the NV centers. Each nanopillar was identified by its fixed grid position and number, allowing one-to-one Hahn-echo measurements with ES cells present and subsequently after cell removal. This design is critical for interpreting the measurements quantitatively: intrinsic differences in NV coherence, depth, and orientation can obscure cellular effects when different sensors are compared. By revisiting each identical sensor, the measured change more directly reflects a contribution of the cellular environment, although other changes associated with cell removal cannot be entirely excluded.

For DOX-treated ES cells, Hahn spin-echo measurements revealed substantially shorter NV spin coherence times (T₂) in the presence of cells than after their removal (Figure 3(H)). Measurements performed on four individually identified nanopillars showed an increase in T₂ after cell removal for every sensor. The percentage increases were 62.2%, 17.9%, 78.8%, and 87.6% for nanopillars 1–4, respectively (Figure 3(I)). These results demonstrate that the coherence shortening observed in the presence of DOX-treated cells was associated with the local cellular environment and was at least partially reversible following cell removal. Because the same NV centers were measured before and after cell removal, the observed changes cannot be attributed solely to differences in intrinsic spin coherence between different nanopillars. The recovery of T₂ following removal of DOX-treated cells is consistent with the reduction of additional local spin fluctuations associated with the cellular environment, potentially originating from nearby paramagnetic species, including free radicals.

To determine whether similar coherence changes occurred in undifferentiated cells, we performed corresponding measurements using DOX− ES cells (Figure 3(J)). Unlike the consistent T₂ recovery observed for DOX+ cells, the DOX− measurements exhibited smaller and less systematic changes following cell removal. The percentage changes in T₂ for the four nanopillars were +27.9%, −6.9%, −16.4%, and +10.1%, respectively (Figure 3(K)). The presence of both positive and negative changes indicates that removal of undifferentiated cells did not produce the consistent coherence recovery observed under differentiation-inducing conditions. This contrast between DOX− and DOX+ cells suggests that differentiation induction is associated with enhanced local spin fluctuations that contribute to NV spin decoherence.

These paired measurements illustrate the central benefit of the substrate-fixed nanopillar architecture. In DOX+ cultures, each of four individually tracked NV sensors showed T₂ recovery after cell removal, whereas the DOX− controls showed no consistent direction of change. Because the comparison is made within the same sensor rather than between different nanodiamonds, intrinsic sensor-to-sensor variability is substantially reduced. The NV response thus provides a repeatable, label-free readout of local spin fluctuations associated with the cellular environment. The measurements do not, by themselves, chemically identify the fluctuating paramagnetic species or establish an absolute intracellular radical concentration.

Taken together, the oxidation-sensitive fluorescence changes, altered cellular morphology, and paired NV coherence measurements indicate that DOX-induced differentiation is accompanied by changes in the local magnetic environment. The consistent T₂ recovery upon removal of DOX-treated cells is compatible with enhanced spin fluctuations from nearby paramagnetic species, including possible radical contributions, while remaining nonspecific to ROS chemistry. This is a distinct biological application from monitoring functional responses of already differentiated cells: it follows a pluripotent cell population undergoing a developmental transition. The quantitative advantage of identical-sensor comparisons provides the experimental basis for expressing these changes numerically through independent TEMPOL reference measurements

### Quantitative Analysis of Radical-Associated Spin fluctuations Using TEMPOL Reference Measurements

To relate intracellular spin-coherence changes to a known paramagnetic reference, we investigated the stable nitroxide radical TEMPOL using the same type of diamond nanopillar NV sensor. Hahn spin-echo measurements were performed before and after exposure to TEMPOL solutions at 100 nM and 1 µM (Figure 4(A)). The measured T₂ decreased with increasing TEMPOL concentration, reaching approximately 1.8 µs at 100 nM and 0.67 µs at 1 µM. In contrast to the spatially heterogeneous coherence changes observed during stem-cell differentiation, TEMPOL exposure produced a more uniform, concentration-dependent reduction in T₂ across the measured nanopillars. These reference responses provide the starting point for expressing cellular magnetic responses as numerical, TEMPOL-equivalent quantities rather than merely relative changes in coherence.

**Figure 4.**
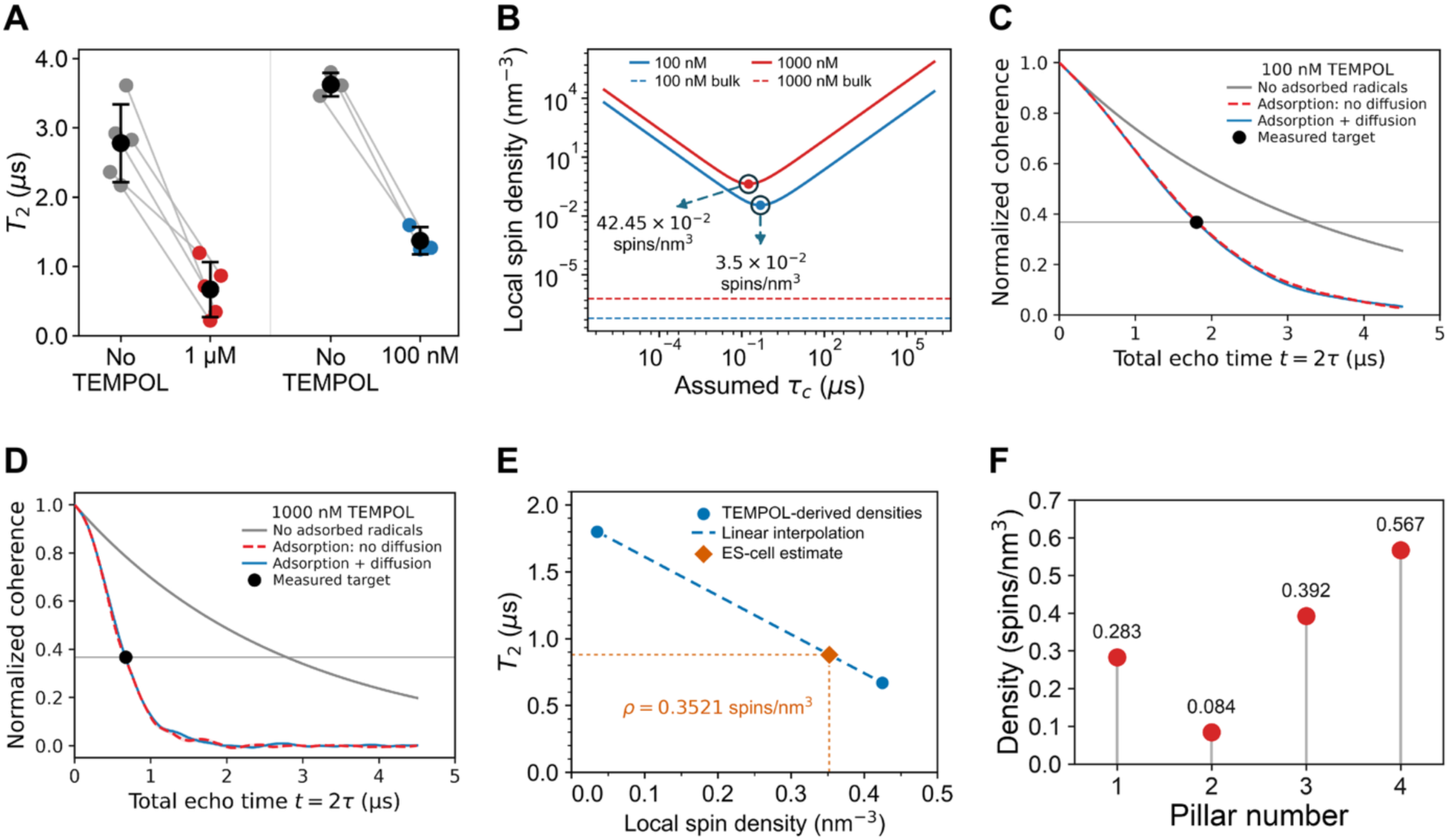
Quantitative analysis of local paramagnetic responses using TEMPOL reference measurements and magnetic-noise modeling. (A) Hahn spin-echo coherence times (T₂) measured using diamond nanopillar NV sensors before and after exposure to 100 nM and 1 µM TEMPOL solutions. (B) Minimum effective local spin densities estimated from experimental T₂ values as a function of the assumed magnetic-field correlation time using a Gaussian magnetic-noise model. (C, D) Kinetic Monte Carlo simulations of Hahn spin-echo coherence for 100 nM (C) and 1 µM (D) TEMPOL, comparing simulated coherence decay under different surface-associated radical conditions with experimental reference values. (E) Two-point linear calibration relating experimentally measured T₂ values to model-derived TEMPOL-equivalent local spin densities, illustrated using a representative measurement from DOX-treated ES cells. (F) TEMPOL-equivalent local spin densities estimated from T₂ measurements at four individually identified NV nanopillar sites in contact with DOX-treated ES cells.

To quantitatively interpret the coherence shortening, we employed a Gaussian magnetic-noise model with exponentially correlated magnetic-field fluctuations. In this model, nearby paramagnetic radicals generate fluctuating dipolar magnetic fields that contribute to NV spin decoherence. The resulting coherence decay depends on the local spin density, dipolar coupling geometry, and magnetic-field correlation time. The effective local spin density was estimated using

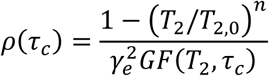

where ρ is the local radical spin density, *τ_c_* is the magnetic-field correlation time, *γ_e_* is the electron gyromagnetic ratio, and *G* represents the dipolar magnetic-field variance per unit spin density calculated for the assumed radical distribution surrounding the diamond nanopillar. *T*_2,0_ denotes the reference coherence time in the absence of the radical-induced contribution, and n is the decay exponent used in the coherence model. The time dependence of the magnetic-noise contribution to Hahn spin-echo decoherence was described by

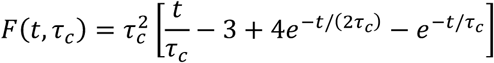

where *t* is the total Hahn-echo evolution time. Because the inferred spin density depends on the magnetic-field correlation time, the minimum local spin density required to reproduce each experimental T₂ value was determined by minimizing ρ(*τ_c_*) over the assumed correlation time.

The resulting minimum effective local spin densities were approximately 3.50 × 10⁻² spins/nm³ for 100 nM TEMPOL and 4.245 × 10⁻¹ spins/nm³ for 1 µM TEMPOL (Figure 4(B)). These values substantially exceeded the corresponding uniform bulk spin densities of 6.022 × 10⁻⁸ and 6.022 × 10⁻⁷ spins/nm³, respectively. This discrepancy indicates that a homogeneous radical distribution at the nominal bulk concentrations cannot readily account for the experimentally observed coherence shortening under the assumed magnetic-noise model. One possible explanation is local enrichment of paramagnetic radicals near the diamond surface, where their magnetic coupling to shallow NV centers is substantially enhanced. Adsorption-mediated enhancement of NV sensing responses has previously been reported for Gd³⁺ ions on carboxylated nanodiamonds by Brown et al. [29]. However, the effective densities inferred from the Gaussian noise model should not be interpreted as direct measurements of the actual molecular concentration or surface coverage, because they depend on the assumed spatial distribution and spin fluctuation dynamics.

To further examine whether surface-associated radical dynamics could account for the observed NV spin decoherence, we performed kinetic Monte Carlo simulations incorporating TEMPOL adsorption, desorption, surface diffusion, and spin reorientation (Figures 4(C) and 4(D)). The simulations explicitly modeled stochastic radical dynamics, with the surface spin populations and characteristic fluctuation times constrained by the preceding Gaussian magnetic-noise analysis. This complementary approach allowed us to evaluate whether the inferred magnetic-noise conditions could reproduce the experimentally observed coherence decay through physically plausible surface-associated radical dynamics. The simulations considered a circular diamond surface region with a radius of 40 nm and an NV center located 10 nm beneath the surface, consistent with the experimentally measured NV depth of 10.5 ± 2.8 nm reported for comparable 3 keV nitrogen implantation conditions [28]. The initial surface coverage was determined using a Langmuir adsorption model, followed by stochastic adsorption, desorption, diffusion, and spin-reorientation events. The resulting fluctuating dipolar magnetic fields were used to calculate the Hahn-echo coherence by averaging over simulated trajectories. Identical kinetic parameters were applied to both TEMPOL concentrations.

At 100 nM TEMPOL, the simulation without adsorbed radicals yielded a baseline coherence time of approximately 3.29 µs, whereas the adsorption-inclusive simulations reproduced a substantially faster coherence decay approaching the experimentally observed T₂ of 1.8 µs (Figure 4(C); Figure S3). Surface diffusion produced a relatively modest additional contribution to coherence shortening, particularly at longer evolution times (Figure S1). At 1 µM TEMPOL, the same kinetic framework predicted substantially faster coherence decay, approaching the experimentally observed T₂ of 0.67 µs near the 1/e coherence level (Figure 4(D); Figure S3). These results indicate that surface-associated radical populations and their stochastic dynamics can account for coherence shortening comparable to the experimental observations at both TEMPOL concentrations under a common set of assumed kinetic parameters. Although these simulations do not independently establish TEMPOL adsorption on the diamond surface or uniquely determine its kinetics, the consistency between the simulated and experimental coherence decays supports surface-associated radical dynamics as a physically plausible explanation for the observed NV spin decoherence under the assumed model conditions. Additional parameter-sensitivity and field-statistics analyses are presented in Figures S1 and S2, while geometry sensitivity is examined in Figure S4.

We next applied the TEMPOL reference measurements to the local spin fluctuations associated with DOX-induced ES-cell differentiation. The minimum effective spin densities obtained from the Gaussian noise model were paired with the corresponding experimental T₂ values to form a two-point reference relationship (Figure 4(E)). Interpolation using a representative T₂ of 0.88 µs in DOX-treated ES cells yielded a TEMPOL-equivalent local spin density of approximately 0.352 spins/nm³. This converts a measured quantum-coherence response into an explicit numerical estimate of effective paramagnetic spin density. However, the value is model-dependent and cannot be interpreted as an absolute ROS concentration, because the spatial distributions, spin dynamics, and NV–spin coupling may differ between the TEMPOL reference solutions and cellular environments.

The fixed array further allowed quantitative comparisons among four individually identified NV sensing sites in DOX-treated ES-cell cultures. Their model-dependent TEMPOL-equivalent local spin densities were estimated as 0.283, 0.084, 0.392, and 0.567 spins/nm³, respectively (Figure 4(F)). The maximum-to-minimum ratio of approximately 6.75 indicates substantial heterogeneity in the inferred local paramagnetic response. The TEMPOL-equivalent spin densities provide a quantitative basis for comparing spatial variations in local paramagnetic responses, enabling site-specific analysis of cellular environments using the fixed NV nanopillar platform.

In summary, the TEMPOL measurements and complementary models enable numerical interpretation of intracellular NV decoherence while keeping the limitations of the inference explicit. The Gaussian magnetic-noise model provides minimum effective local spin-density estimates, and complementary kinetic Monte Carlo simulations support the physical plausibility of surface-associated radical dynamics under the assumed model conditions for the enhanced TEMPOL response. Combined with same-sensor measurements, the resulting model-dependent TEMPOL-equivalent densities reveal spatial differences in the magnetic environment of differentiating ES cells. This proof of concept extends intracellular quantum sensing from measurements of functional changes in mature cell populations toward quantitative readout of physicochemical changes accompanying pluripotent stem-cell differentiation. More broadly, such measurements may help connect cellular-scale magnetic activity to studies of development, tissue function, and future regenerative-medicine applications.

## Conclusion

We established a substrate-fixed diamond nanopillar quantum sensing platform that enables quantitative, label-free investigation of local magnetic environments in living cells through one-to-one measurements using the same nitrogen-vacancy (NV) sensors. By comparing the spin coherence of identical NV centers in the presence and absence of cells, this approach minimizes sensor-to-sensor variability and provides a reliable basis for distinguishing cellular contributions to spin decoherence. Cellular compatibility assays supported the preservation of cellular functions, while coherent quantum measurements demonstrated approximately 10% ODMR contrast, 20% Rabi contrast, and a Hahn-echo T₂ of approximately 3 µs inside living cells. CPMG dynamical decoupling further extended intracellular T₂ to 25.8 ± 4.90 µs, demonstrating the ability of the fixed nanopillar architecture to maintain prolonged quantum coherence in cellular environments.

We applied this platform to investigate the differentiation of embryonic stem (ES) cells toward the trophectoderm lineage following doxycycline-mediated Oct3/4 repression. All four individually tracked nanopillars exhibited recovery of spin coherence after removal of differentiation-induced cells, whereas undifferentiated controls showed no consistent recovery. These results, together with complementary oxidation-sensitive fluorescence measurements, indicate differentiation-associated changes in the local magnetic environment. Using the stable radical TEMPOL as a reference, together with Gaussian magnetic-noise analysis and complementary kinetic Monte Carlo simulations, we quantified spatial variations in local magnetic responses as model-dependent TEMPOL-equivalent spin densities of 0.084–0.567 spins/nm³. These values provide a numerical measure of local paramagnetic activity rather than an absolute intracellular ROS concentration.

By extending intracellular quantum sensing from functional responses of differentiated cells to the differentiation of pluripotent stem cells, this work establishes a quantitative approach for investigating physicochemical changes associated with developmental processes. The combination of stable, individually addressable quantum sensors, direct one-to-one comparisons, and reference-based quantification opens new opportunities to investigate cellular processes underlying tissue formation and regeneration, demonstrating the potential of quantum sensing for future applications in regenerative medicine.

## Methods

### Diamond nanopillar fabrication

NV-containing diamond nanopillar arrays were fabricated from single-crystalline diamond substrates with a (100) surface orientation and a residual nitrogen concentration below 10 ppb. The substrates had lateral dimensions of 2 × 2 mm², a thickness of 40 ± 10 µm with a thickness variation of less than 1 µm, and polished surfaces with a root-mean-square (RMS) roughness below 1 nm. Near-surface NV centers were formed by implantation of ^¹⁴^N ions at an energy of 3 keV, with the implantation dose selected to produce a high NV-center areal density exceeding 1000 NVs/µm². An NV depth of 10 nm was assumed in the calculations based on the experimentally measured mean depth of 10.5 ± 2.8 nm reported by Pham et al. for NV centers formed by 3 keV ¹⁵N implantation [28], rather than on a direct depth measurement of the present ¹⁴N-implanted devices.

Diamond nanopillar arrays were fabricated using electron-beam lithography and oxygen-based inductively coupled plasma (ICP) etching. Briefly, electron-beam lithography and metallization were used to define nanoscale etching masks on the diamond surface. The exposed diamond was subsequently removed by oxygen-plasma etching, leaving nanopillars beneath the masked regions. The nanopillar geometry and sidewall profile were controlled through the mask dimensions and ICP etching conditions. The resulting nanopillars had a diameter of 300 nm, a height of 3.2 µm, and a center-to-center spacing of 15 µm.

### Optical, microwave, and NV spin measurements

Measurements were performed using an Olympus IX73 inverted microscope, with the excitation laser directed to the objective through a dichroic mirror. A high-power laser (model: Verdi G5) was gated using an acousto-optic modulator (Gooch & Housego, 3250-220) driven by an RF driver (3910-XX). Collected fluorescence was directed to a pinhole-coupled avalanche photodiode (SPCM-AQRH-14-FC-ND), or a color CCD camera (model E3CMOS). Temperature was controlled using an incubator mounted on a piezoelectric stage (P-545.3C8S) with thermocouple feedback.

Microwaves were delivered to the NV centers through a 20 µm-diameter copper wire connected to sputtered Ti/Cu/Au electrodes on a glass coverslip. Signals were generated using an N5182A RF signal generator and amplified using an ZHL-50 W-63+. Pulse sequences were controlled using a DTG5274 pulse generator, with microwave pulses gated by a Mini-Circuits ZASWA-2-50DRA+ RF switch.

NV spin properties were characterized using ODMR, Rabi oscillation, spin-echo, and CPMG measurements. ODMR spectra were obtained by sweeping the microwave frequency while monitoring fluorescence, with the contrast calculated as

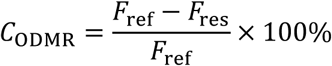

Optically detected magnetic resonance (ODMR) spectroscopy was performed by sweeping the microwave frequency and monitoring NV fluorescence to identify the spin resonance frequencies. Rabi measurements were performed by varying the resonant microwave-pulse duration to determine the Rabi frequency and calibrate the 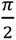 and *π* pulse lengths. Spin-echo measurements used a 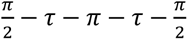 sequence, and the coherence decay was fitted using

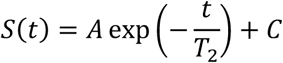

where *t* = 2*τ*. CPMG measurements were performed using multiple refocusing pulses to evaluate dynamical-decoupling-enhanced coherence. ODMR contrast, Rabi contrast, and spin-echo coherence times were characterized across five devices.

ODMR, Rabi, and spin-coherence measurements were performed on individually addressed nanopillars associated with ES cells. Cells were maintained at a temperature of 37^0^ C during acquisition. Spin-echo measurements were compared among cell-free nanopillars, nanopillars associated with DOX− cells, and nanopillars associated with DOX+ cells. Additional measurements compared nanopillars classified as intracellular and extracellular in DOX+ cultures.

For each measurement site, the microwave resonance frequency was identified using ODMR spectroscopy, and the π/2 and π pulse durations were calibrated using Rabi oscillation measurements. Echo decays were acquired based on π/2 and π pulse durations and fitted using the exponential function described above.

### ES-cell culture and differentiation

ZHBTc4 mouse embryonic stem (ES) cells carrying a tetracycline-regulatable Oct3/4 transgene were maintained in Glasgow minimum essential medium (GMEM) supplemented with 10% fetal bovine serum (FBS), 1 mM sodium pyruvate, 0.1 mM nonessential amino acids, 0.1 mM 2-mercaptoethanol, and 1000 U/mL leukemia inhibitory factor (LIF) at 37 °C in a humidified atmosphere containing 5% CO₂. Cells were seeded onto diamond nanopillar arrays and cultured for 72 hrs before measurements. Trophectoderm differentiation was induced by repression of Oct3/4 using the Tet-Off system. Cells were treated with doxycycline (DOX) and addition of DOX represses expression of the tetracycline-regulated Oct3/4 transgene, promoting differentiation toward the trophectoderm lineage. Cells maintained without DOX served as the undifferentiated DOX− control.

### Cellular compatibility assessments

Cellular compatibility was evaluated through fluorescence assays of metabolic activity, apoptosis, DNA synthesis, and plasma membrane integrity. Metabolic activity was assessed using the alamarBlue Cell Viability Reagent (Invitrogen DAL1100). Apoptotic activity was assessed using the CellEvent Caspase-3/7 Green Detection Reagent (Thermo Fisher Scientific, C10723) and green nuclear fluorescence was examined as an indicator of caspase-3/7 activation. Positive controls were prepared using apoptosis-inducing treatment. DNA synthesis was assessed using the Click-iT EdU Cell Proliferation Kit (Thermo Fisher Scientific, C10337). Cells were incubated with EdU, followed by fixation, permeabilization, and fluorescent labeling according to the manufacturer’s protocol. Nuclei were counterstained with Hoechst nucleic acid stain (Dojndo 346-07951). Blue fluorescence identified nuclei, whereas green fluorescence identified cells that incorporated EdU into newly synthesized DNA during the labeling period. Plasma membrane morphology and integrity were assessed using CellMask Plasma Membrane Stain (Invitrogen C37608), NucBlue Live reagent, and NucGreen Dead reagent (Invitrogen R37609). CellMask delineated the plasma membrane, NucBlue labeled nuclei, and nuclear uptake of NucGreen indicated compromised membrane integrity. Intracellular oxidative activity during differentiation was assessed using CM-H₂DCFDA (Invitrogen C6827). DOX− and DOX+ ES cells were incubated, and green fluorescence was acquired using identical imaging settings for both groups. The fluorescence response was interpreted as an indicator of intracellular oxidative activity. Fluorescence images were acquired using confocal laser scanning microscopy (LSM). GFP-labeled actin filaments and Hoechst-stained nuclei were imaged in DOX− and DOX+ cells to assess cell morphology and cytoskeletal organization.

### TEMPOL reference measurements and calibration

TEMPOL was dissolved in ultrapure water to prepare solutions at concentrations of 100 nM and 1000 nM. Measurements were performed using the optical and microwave conditions described above. Each concentration was referenced to its corresponding TEMPOL-free baseline. Spin-echo decays were fitted to obtain T2 values for the reference and TEMPOL-containing conditions.

A two-point linear calibration was constructed using the experimentally measured T₂ values and corresponding model-derived effective local spin densities for the 100 nM and 1 µM TEMPOL reference conditions. Cellular T₂ values were then converted into model-dependent TEMPOL-equivalent spin-density estimates using this relationship. Applying this relationship to cellular measurements assumed comparable background decoherence, NV depths, and sensing geometries, together with similar effects of surface adsorption and radical fluctuation dynamics on the sensing response. The resulting estimates were therefore conditional on these assumptions and represented model-dependent TEMPOL-equivalent local spin densities rather than direct measurements of total intracellular ROS concentration.

### Simulation of surface-associated TEMPOL

TEMPOL adsorption and its contribution to NV Hahn-echo dephasing were investigated using continuous-time kinetic Monte Carlo simulations. The simulated surface consisted of a circular patch of radius 40 nm on the nanopillar top surface. The NV center was positioned 10 nm below the patch center, with its quantization axis normal to the surface. Adsorbed radical spins were located 0.37 nm above the surface, and each lattice site accommodated at most one radical. The surface exchanged radicals with a well-mixed reservoir maintained at a fixed TEMPOL concentration of 100 or 1000 nM. Initial site occupancy was sampled independently from Langmuir equilibrium,

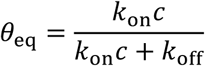

where *c* is the reservoir concentration in molar units, *k*_on_ is the adsorption rate constant, and *k*_off_ is the desorption rate. Four stochastic processes were included: adsorption, desorption, surface hopping, and phenomenological spin reorientation. The assumed parameters were *k*_on_ = 10^11^ M^−1^ s^−1^, *k*_off_ = 10^5^ s^−1^ and *k*_rot_ = 10^6^ s^−1^ Simulations with *D*_s_ = 0 and *D*_s_ = 10^−12^ m^2^ s^−1^ were compared using identical kinetic parameters at both concentrations. A control with *k*_on_ = 0 represent the absence of adsorbed radicals.

Radical magnetic moments were represented by isotropic classical spin vectors whose second moments matched those of unpolarized spin ^1^ particles. The magnetic-field component parallel to the NV axis was 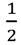 calculated by summing the dipolar contributions from all occupied sites:

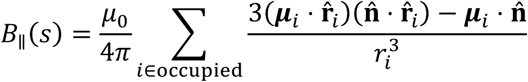

Where ***μ****_i_* is the radical magnetic moment, *r_i_*is its distance from the NV center, **r**^*_i_* is the corresponding unit vector, and *n* is the NV quantization axis. The field was constant between stochastic events and was integrated directly over each event interval. For trajectory *j* the accumulated Hahn-echo phase was

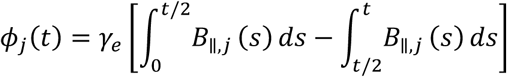

where *t* = 2*τ* is the total free-evolution time and *γ_e_* is the electron gyromagnetic ratio in angular-frequency units. Ideal instantaneous refocusing pulses were assumed. The simulated coherence was obtained as

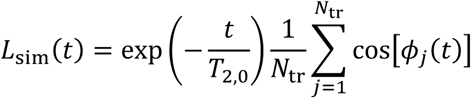

Here, *N*_tr_ = 32 independent trajectories were averaged for each condition, and *T*_2,0_ = 3.0 *μ*s represented an independent background decay, with the dry-device baseline assumed to apply in solution. Coherence was evaluated at 91 equally spaced times from 0 to 4.5 µs. Pointwise Monte Carlo standard errors were calculated across trajectories, and the simulated *T*_2_ was determined by linear interpolation at the first crossing of *L*_sim_(*T*_2_) = *e*^−1^.

The measured coherence times were used as comparison targets without fitting the kinetic parameters. No single magnetic-field correlation time was imposed. Reservoir depletion, freely dissolved radical contributions, nanopillar sidewalls, and explicit radical–radical spin dynamics were excluded. The simulations therefore evaluated adsorption as a possible dephasing mechanism under the specified assumptions.

## References

1. Dröge, W. Free radicals in the physiological control of cell function. Physiol. Rev. 82, 47–95 (2002).

2. Jang, Y. Y. & Sharkis, S. J. A low level of reactive oxygen species selects for primitive hematopoietic stem cells that may reside in the low-oxygenic niche. Blood 110, 3056–3063 (2007).

3. Sauer, H. et al. Role of reactive oxygen species and phosphatidylinositol 3-kinase in cardiomyocyte differentiation of embryonic stem cells. FEBS Lett. 476, 218–223 (2000).

4. Kobayashi, C. I. & Suda, T. Regulation of reactive oxygen species in stem cells and cancer stem cells. J. Cell. Physiol. 227, 421–430 (2012).

5. Ji, A. R. et al. Reactive oxygen species enhance differentiation of human embryonic stem cells into mesendodermal lineage. Exp. Mol. Med. 42, 175–186 (2010).

6. Sies, H. et al. Defining roles of specific reactive oxygen species (ROS) in cell biology and physiology. Nat. Rev. Mol. Cell Biol. 23, 499–515 (2022).

7. Zhang, Y. et al. Methods for the detection of reactive oxygen species. Anal. Methods 10, 4625–4638 (2018).

8. Chen, X. et al. Fluorescent and luminescent probes for detection of reactive oxygen and nitrogen species. Chem. Soc. Rev. 40, 4783–4804 (2011).

9. Chen, X. et al. Recent progress in the development of fluorescent, luminescent and colorimetric probes for detection of reactive oxygen and nitrogen species. Chem. Soc. Rev. 45, 2976–3016 (2016).

10. Waters, J. C. Accuracy and precision in quantitative fluorescence microscopy. J. Cell Biol. 185, 1135– 1148 (2009)

11. A Sigaeva et al. Diamond-based nanoscale quantum relaxometry for sensing free radical production in cells Small 18 (44), 2105750

12. L Nie et al, Quantum sensing of free radicals in primary human dendritic cells Nano Lett 22 (4), 1818–1825

13. R Sharmin et al Fluorescent nanodiamonds for detecting free-radical generation in real time during shear stress in human umbilical vein endothelial cells ACS sensors 6 (12), 4349

14. K Wu et al, Applying NV center-based quantum sensing to study intracellular free radical response upon viral infections Redox biology 52, 102279

15. L Wu et al, Diamond relaxometry as a tool to investigate the free radical dialogue between macrophages and bacteria ACS nano 17 (2), 1100–1111

16. N Norouzi et al, Relaxometry for detecting free radical generation during Bacteria’s response to antibiotics Carbon 199, 444–452

17. F Perona Martínez et al. Nanodiamond relaxometry-based detection of free-radical species when produced in chemical reactions in biologically relevant conditions. ACS sensors 5 (12), 3862–3869

18. S Fan et al, Quantum sensing of free radical generation in mitochondria of single heart muscle cells during hypoxia and reoxygenation ACS nano 18 (4), 2982–2991

19. Mzyk, A. et al. Quantum sensing to monitor changes in free radical generation by intracellular vesicles of polarized macrophages. Acta Biomater. 199, 315–323 (2025). 10.1016/j.actbio.2025.04.024.

20. Mzyk, A. et al. Nanodiamond-Based Quantum Sensing Reveals Changes in Endo-Lysosomal Activity of Stimulated Cardiac Fibroblasts in a Cellular Fibrosis Model. Adv. Funct. Mater. 36, e28248 (2026). 10.1002/adfm.202528248.

21. Fujisaku, T. et al. pH nanosensor using electronic spins in diamond. ACS Nano 13, 11726–11732 (2019).

22. Oshimi, K. et al. Bright quantum-grade fluorescent nanodiamonds. ACS Nano 18, 35202–35213 (2024). 10.1021/acsnano.4c03424.

23. Gu, Q. et al. Simultaneous nanorheometry and nanothermometry using intracellular diamond quantum sensors. ACS Nano 17, 20034–20042 (2023).

24. Niora, M. et al. Bio-sensing with NV-centers in diamond. J. Phys. Photonics 8, 012007 (2026).

25. Losero, E. et al. Neuronal growth on high-aspect-ratio diamond nanopillar arrays for biosensing applications. Sci. Rep. 13, 5909 (2023).

26. Niwa, H. et al. Interaction between Oct3/4 and Cdx2 determines trophectoderm differentiation. Cell 123, 917–929 (2005).

27. Niwa, H.; Miyazaki, J.; Smith, A. G. Quantitative expression of Oct-3/4 defines differentiation, dedifferentiation or self-renewal of ES cells. Nature Genetics 24, 372–376 (2000).

28. Pham, L. M. et al. NMR technique for determining the depth of shallow nitrogen-vacancy centers in diamond. Phys. Rev. B 93, 045425 (2016). 10.1103/PhysRevB.93.045425.

29. Brown, E. K. et al. Quantum sensing of paramagnetic analytes by nanodiamonds in levitated microdroplets and aqueous solutions. Chemical Science 16, 19328–19341 (2025).

