## Supplementary material for "Quantification of Spin Fluctuations during Stem Cell Differentiation Using Diamond Nanopillars": SI

### Contents

**Figure S1.** Sensitivity of simulated Hahn-echo coherence to kinetic parameters ··· S2

**Figure S2.** Compensated desorption-rate sensitivity ··· S3

**Figure S3.** Monte Carlo convergence and uncertainty analysis ··· S4

**Figure S4.** Sensitivity of inferred spin density to NV depth ··· S5

**Figure S5.** TEMPOL calibration and site-specific density estimates ··· S6

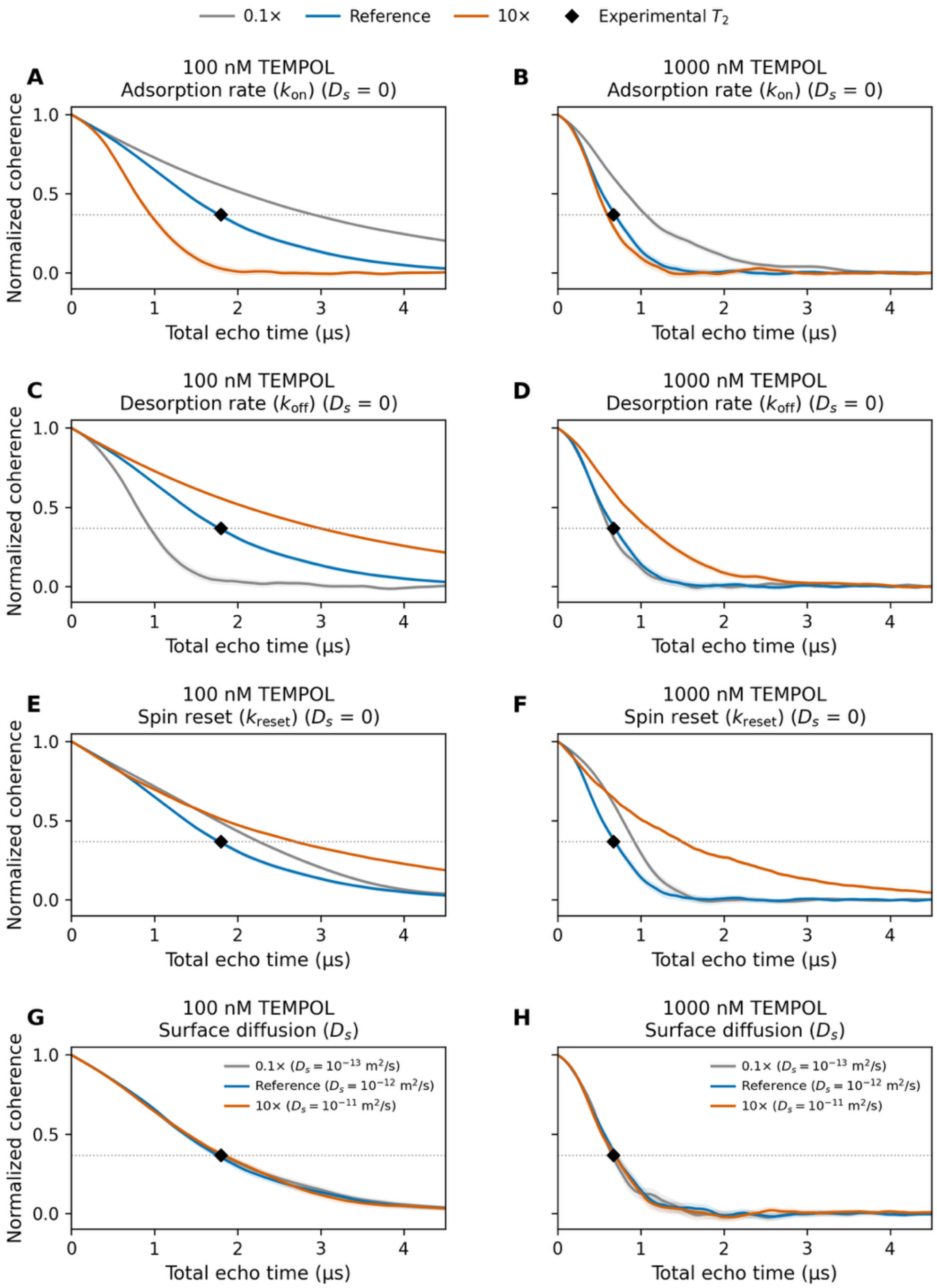


**Figure S1.** Sensitivity of simulated Hahn-echo coherence to (A, B) adsorption rate $k_{on}$, (C, D) desorption rate $k_o$ff, (E, F) spin-reset rate $k_{reset}$, and (G, H) surface diffusion coefficient $D_s$ for 100 nM (left) and 1 μM (right) TEMPOL. Parameters were varied individually to 0.1× (gray), 1× (blue), and 10× (orange) their reference values.

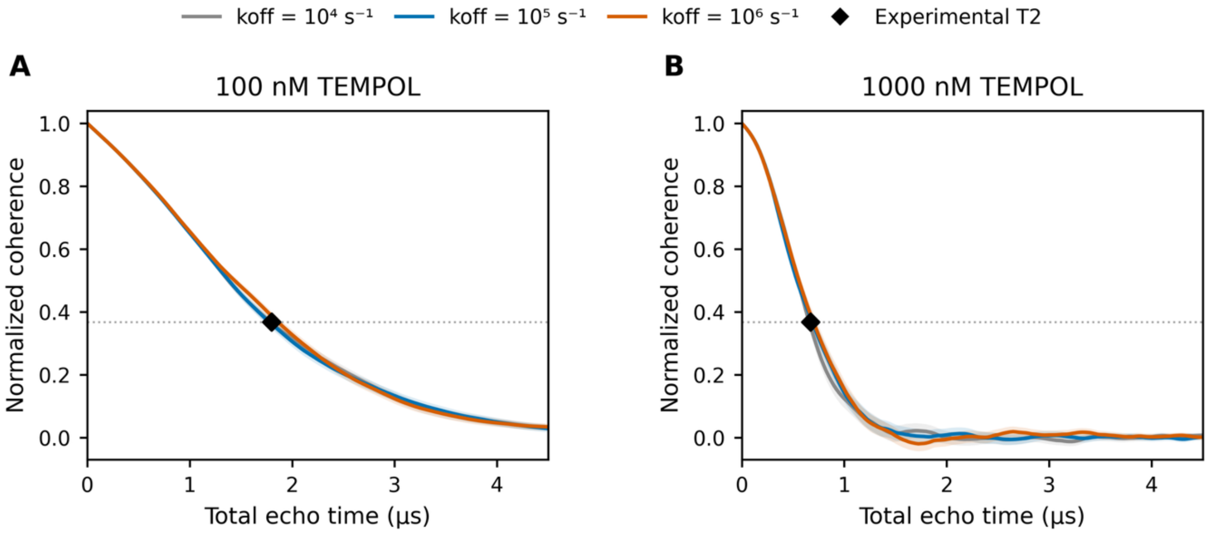


**Figure S2.** Hahn-echo coherence at (A) 100 nM and (B) 1 µM TEMPOL for $k_{o}ff = 10^4$, $10^5$, and $10^6$ $s^{-1}$. For each $k_{o}ff$, $k_{on}$ and $k_{reset}$ were recalculated to preserve surface occupancy and the target field-correlation time at $D_s = 0$.

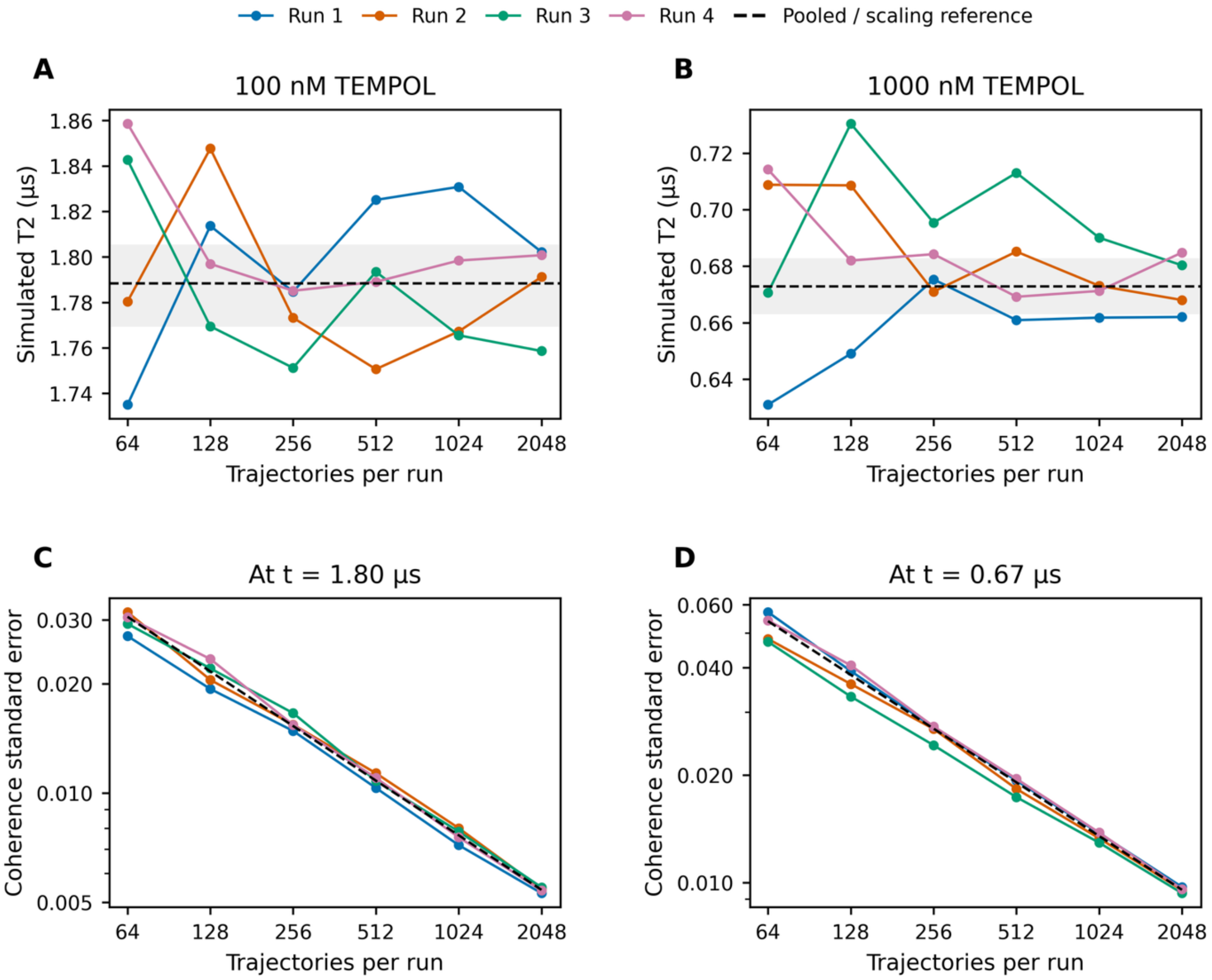


**Figure S3.** Four independently seeded runs at each TEMPOL concentration, using nested subsets of 64 to 2,048 trajectories per run. (A, B) Simulated $T_2$ from the first 1/e crossing; dashed lines and gray bands show the pooled estimate and 95% bootstrap interval from 8,192 trajectories per concentration. (C, D) Standard error of coherence at the experimental $T_2$; dashed lines show $1/\sqrt{N}$ scaling. Reference rates were used with $D_s = 0$.

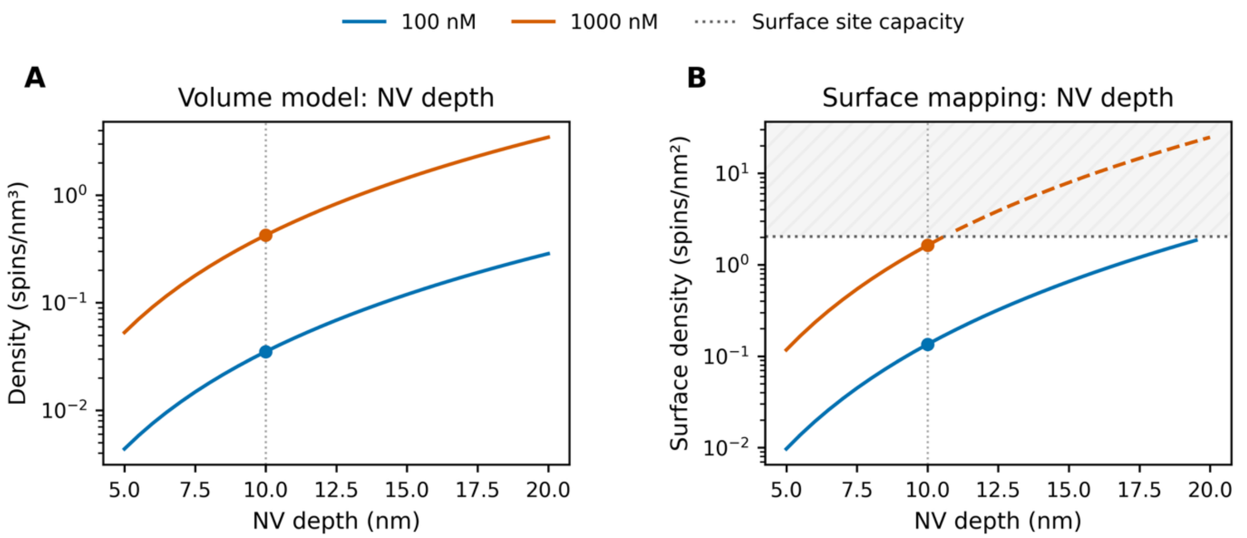


**Figure S4.** Sensitivity of model-inferred TEMPOL spin density to NV depth. (A) Required volumetric spin density and (B) variance-equivalent surface spin density as functions of NV depth. The target $T_2$, corresponding baseline coherence time, and selected correlation time ($\tau c$) were held constant. Dots indicate the reference NV depth of 10 nm. In (B), the horizontal dotted line represents the maximum capacity of the 0.7 nm surface lattice, while the hatched region and dashed curve segments indicate densities exceeding this limit.

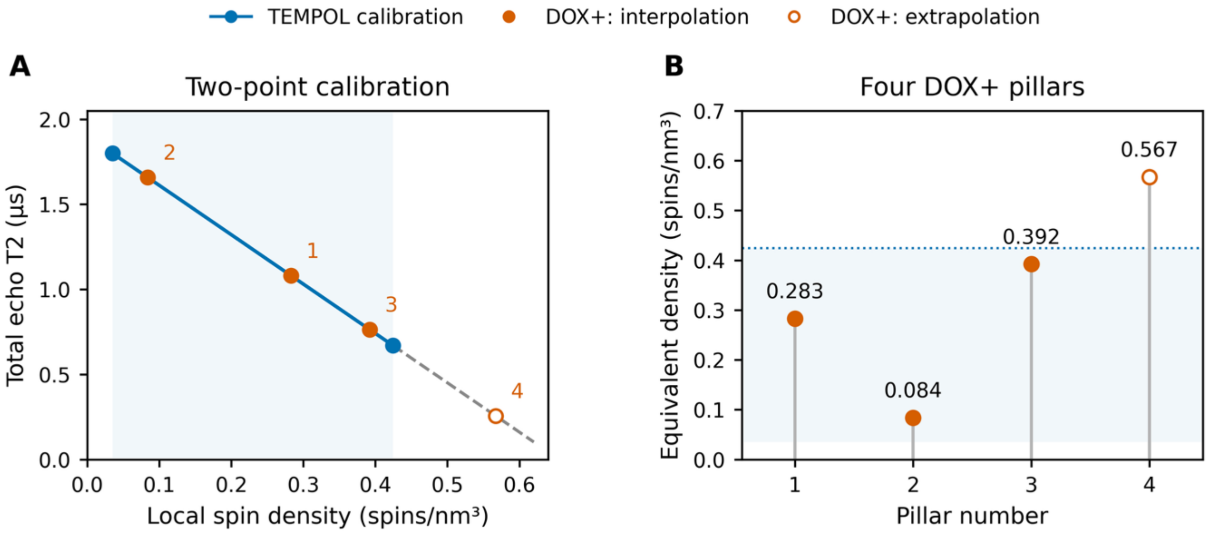


**Figure S5.** (A) Two-point linear calibration using densities of 0.0350 and 0.4245 spins/nm³ at $T_2 = 1.80$ and 0.67 µs. Numbered points show the four DOX+ pillars. (B) Corresponding TEMPOL-equivalent density estimates. Shading marks the calibration range; the dashed extension and open marker identify extrapolation for pillar 4. The supplied time multiplier of 2.0 was retained. No experimental uncertainty estimates were supplied.
